# Walking speed can be modulated on an adaptive split-belt treadmill

**DOI:** 10.1101/2025.06.03.657157

**Authors:** Rucha Kulkarni, Barry Bodt, Jill S. Higginson

## Abstract

Adaptive treadmills (ATMs) that change speed based on the users’ gait mechanics can allow for healthy stride-to-stride variability observed during overground walking, while maintaining the benefits of traditional treadmill training for gait rehabilitation. To enable unilateral targeting of propulsion, we developed an adaptive split-belt treadmill (sATM) that updates the speed of each belt of the treadmill based on the user’s propulsion and position. The present study demonstrated equivalence of the sATM to an existing tied-belt ATM that updates the speed of both belts based on the user’s propulsion, step length, and position. Fourteen young, healthy participants completed five trials at their comfortable walking speed on the tied ATM and sATM with unilateral and bilateral modifications. We hypothesized that propulsion and belt speed on the novel sATM would be equivalent to the tied ATM, and that the sATM could be modified to encourage higher propulsion unilaterally while maintaining walking speed. Participants maintained similar propulsion and belt speeds between treadmills and belts for bilateral modifications. While some participants increased unilateral propulsion to maintain similar walking speeds for the unilateral modification, others showed no change in propulsion with slight differences in walking speed. Our results suggest that the response to unilateral modifications was user-specific, and the sATM could be tuned individually to allow unilateral control strategies. The present study demonstrates that walking speed can be modulated on an adaptive split-belt treadmill.

## 1. Introduction

Walking speed is often used as a marker of healthy gait (Fritz and Lusardi, 2009) and community ambulation in older adults and neurological conditions like stroke (Dickstein, 2008; Fritz and Lusardi, 2009). It is heavily influenced by propulsion (push-off), which can be defined as the positive work performed by the limb to accelerate the body forward (Hsiao *et al*., 2015; Awad *et al*., 2020). For people with post-stroke hemiparesis, the weakened muscles on one side of the body reduce the propulsion forces generated by the paretic (affected) limb, which contributes to a decreased walking speed and affects community ambulation (Patterson *et al*., 2007).

Treadmill training is commonly used in gait rehabilitation for stroke survivors as it enables high repetition of movement in a safe and controlled environment (Dickstein, 2008; Duncan *et al*., 2011). However, conventional treadmills operate at fixed speeds which limits any instantaneous changes in speed and propulsion that we might observe in overground walking. Previous rehabilitation paradigms have been limited to fixed-speed treadmills which limits stride-to-stride variability that is essential for motor learning (Stergiou and Decker, 2011). Overall, traditional treadmill training shows mixed efficacy and limited improvement in walking speed and propulsion (Gelaw *et al*., 2019).

Split-belt treadmills have been developed such that the walking surface is composed of two belts with speeds set independently. Using this tool, a series of studies has been conducted with the speed of one limb set to twice the other and have shown to improve spatiotemporal gait characteristics in stroke survivors (Reisman *et al*., 2007, 2013). However, the speed differential is controlled by a clinician and may lead to sub-optimal rehabilitation conditions. Furthermore, split-belt treadmills have also been restricted to fixed speeds, which limit stride-to-stride variability (a characteristic of overground walking) and prevent targeting of specific gait kinetics like propulsion. This may play a role in the retention and limited transfer of adapted gait mechanisms from split-belt treadmills to overground walking, which is the ultimate goal of walking rehabilitation (Reisman *et al*., 2009).

To allow for stride-to-stride variability while retaining any benefits of treadmill training, adaptive treadmills (ATMs) that update speed based on the user gait mechanics (user-driven) have been developed (Sloot, Van Der Krogt and Harlaar, 2014; Ray, Knarr and Higginson, 2018; Pariser *et al*., 2022). Early treadmill control schemes involved user position on the treadmill or spatiotemporal gait characteristics, and did not consider the relationship between propulsion and walking speed (Sloot, Van Der Krogt and Harlaar, 2014). Recently developed ATMs update speed based on propulsion, step length and the user’s position on the treadmill relative to the treadmill center (Ray, Knarr and Higginson, 2018; Pariser *et al*., 2022). Studies have shown that healthy individuals walk on an ATM at a self-selected speed closer to overground walking compared to fixed-speed treadmills suggesting that ATMs may be more similar to overground walking (Ray, Knarr and Higginson, 2018; Kempski *et al*., 2019; Donlin *et al*., 2022). However, on average, stroke survivors do not seem to increase their self-selected speed on the ATM as compared to fixed-speed treadmills (Ray, Reisman and Higginson, 2020). Previous work has also demonstrated that healthy individuals can increase bilateral propulsion to maintain a consistent walking speed on these ATMs with increased propulsion demand (increasing the control parameter for propulsion) (Pariser *et al*., 2022). However, to date, these ATMs restrict both limbs to the same speed and thus do not target unilateral gait mechanics. The purpose of this study was to develop and validate an adaptive split-belt treadmill (sATM) to enable preferential targeting of unilateral gait mechanics. The objective of this study was to determine how healthy young adults modulate speed on the novel adaptive split-belt treadmill. We hypothesized that: 1) when the control parameters were equal on both belts on the split-belt treadmill and equal to that of a tied-belt treadmill developed by Pariser et al. (2022), there would be no difference in propulsion and belt speed between treadmills; 2) when the control parameter for propulsion (propulsion demand) was increased equally on both belts, propulsion would increase to maintain belt speed; and 3) when propulsion demand was increased for just one belt on the split-belt treadmill, propulsion on the targeted belt would increase with no difference in belt speed.

## 2. Methods

### 2.1.1 sATM controller

The previous implementation of the adaptive tied-belt (Tied) treadmill updates the speed of both belts based on the current speed, the intermediate speeds due to average propulsion impulse (net anterior impulse) and average step length of both limbs, and the position of the person’s center of mass (COM) relative to the treadmill center (Pariser *et al*., 2022). Intermediate speeds are calculated during swing phase using data from the immediately previous stance phase, and represent belt speed if only the respective metric determined the overall speed (Pariser *et al*., 2022).

For application to the split-belt adaptive treadmill (Split), the updated speed of each belt (*v*_*i*+1_) is calculated independently at each step based on the current speed (*v*_*i*_), the intermediate speed due to propulsion (net anterior-posterior impulse of the respective limb) (*v*_*PI*_), and the position of the person’s center of mass relative to the treadmill center (*COM*_*pos*_) (Fig 1A)

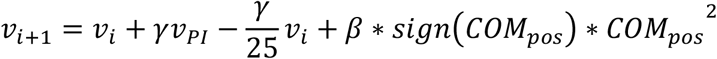

where 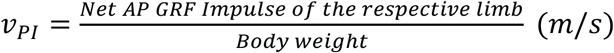 and 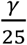 is a tuned value from pilot testing.

**Fig 1.**
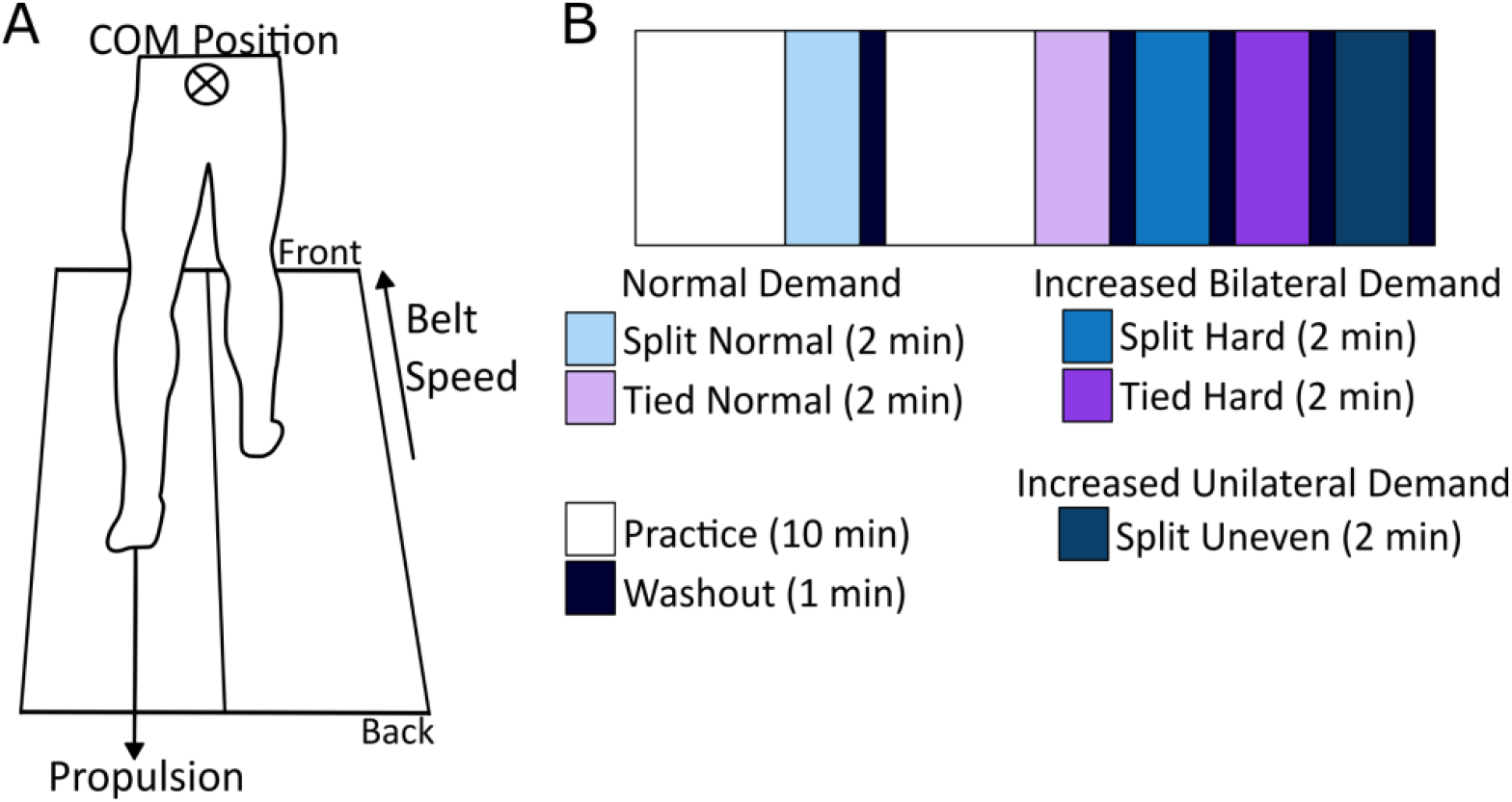
(A) Experimental Set-Up. An illustrative sketch of a participant on the sATM. (B) Experimental Protocol. Participants performed 5 trials on the Tied ATM and Split ATM.

Thus, increasing or decreasing propulsion increases or decreases the overall belt speed respectively. Similarly, moving forward (increasing the center of mass position) relative to the treadmill center increases speed and moving backwards decreases speed.

The peak propulsive force was also calculated in real-time as the maximum anterior ground reaction force (AGRF) per stride. Peak AGRF asymmetry was then calculated as the difference of the contralateral and ipsilateral peak AGRF divided by the sum.

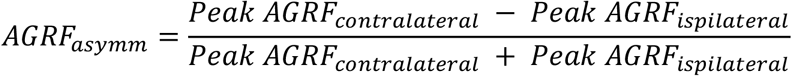

To promote healthy gait symmetry, when peak AGRF asymmetry (*AGRF*_*asymm*_) was less than 5% (Lauzière *et al*., 2014; Herzog *et al*., Feb1989), the updated belt speed was calculated using the intermediate speed due to propulsion (net anterior-posterior impulse) averaged over both limbs:

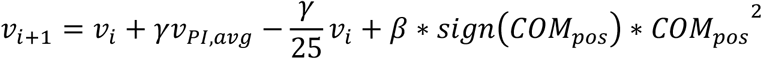

where 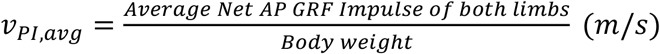.

For peak AGRF asymmetry (*AGRF*_*asymm*_) greater than 5%, a belt speed penalty is imposed based on the level of asymmetry. Essentially, the updated belt speed is decreased by the intermediate speed due to propulsion (net anterior-posterior impulse) averaged over both limbs around this threshold to remove this discontinuity:

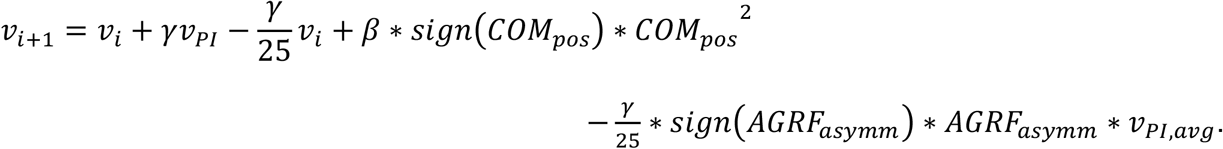

The updated speed was calculated between heel-strike and toe-off and updated at toe-off of each stride. To ensure smooth belt accelerations, the propulsion gain (*γ*) was calculated using an exponential function where dt is the time between belt speed updates (i.e. the time from heel-strike to toe-off for each stride) and the position gain (*β*) was a tuned value from prior work (Ray, Knarr and Higginson, 2018; Pariser *et al*., 2022):

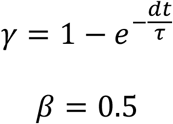

The value of tau (*τ*) was set to either 1.5 or 0.7639 to vary gamma, described more in section 2.1.2. Decreasing tau (*τ*) (increasing gamma (*γ*)) requires higher propulsion at each step to maintain the user’s self-selected walking speed.

### 2.1.2 Data Collection

Fourteen adults (24.79 ± 1.82 years, 1.70 ± 0.08 m, 75.86 ± 14.91 kg, 7 females) participated and self-reported no known neuromuscular injuries or disorders. The University of Delaware Institutional Review Board approved the protocol, and participants gave written informed consent and completed a modified physical activity readiness questionnaire. Participants first performed the 10-meter overground walking test to determine their preferred overground walking speed. Then they performed a series of walking trials on an instrumented split-belt treadmill (Bertec Corp; 2000 Hz) and real-time data were collected using a motion capture data collection system (Qualisys; 100 Hz).

To demonstrate equivalence, we asked participants to walk on the novel adaptive split-belt treadmill (sATM, Split treadmill) and on a tied-belt adaptive treadmill (Tied treadmill) previously developed by Pariser et al. (2022). The propulsion demand (gain) was varied bilaterally (Normal and Hard conditions) to establish equivalence for normal and increased bilateral propulsion demand and unilaterally (Uneven) to understand the effect of increased unilateral propulsion demand. Participants performed 5 walking trials for 2 minutes each – Split Normal, Tied Normal, Tied Hard, Split Hard, and Split Uneven in a pseudo-randomized order (Fig 1B). For the Normal and Hard conditions on the Split treadmill, the propulsive gains of both belts were identical to the Tied Normal and Hard conditions. The Hard conditions (*τ*_*γ*_ = 0.7639, *γ*_*Hard*_) had a higher propulsive gain (*γ*_*Hard*_ = 1.5\**γ*_*Normal*_) than the Normal condition (*τ*_*γ*_ = 1.5, *γ*_*Normal*_) and required higher propulsion to maintain a comfortable speed, as observed by Pariser et al. For the Uneven condition, one belt was defined as the Targeted side and had a higher propulsive gain (*γ*_*Hard*_) than the other side (*γ*_*Normal*_), such that *γ*_*Targeted*_= 1.5 ∗ *γ*_*Non*−*targeted*_. The Targeted side (right or left belt) was randomized for all participants (N=7 right).

Participants were given up to 10 minutes practice for each of the Normal conditions before performing the respective Normal trial. Both Normal trials (Split and Tied) were performed first in random order, followed by the Hard and Uneven trials in random order. Participants were given up to 3 minutes to achieve a comfortable, steady-state speed and the trial began when they gave verbal indication that they had achieved a comfortable speed. The maximum speed of the treadmill was set to 2 m/s. Participants were allowed to lightly touch the handrails for stabilization. After each trial, participants were asked to respond to a survey to assess their perceived: (1) stability, (2) overall comfort, (3) ease of achieving their preferred walking speed, and (4) ease of maintaining their preferred speed. Each metric was measured on a scale of 1–7 where 1 corresponded to very unstable, uncomfortable, or difficult and 7 represented very stable, comfortable, or easy. Participants also performed a washout trial (1 min) of walking on a fixed speed treadmill at 0.1 m/s less than their preferred overground walking speed after each trial.

### 2.1.3 Data Processing and Analysis

Kinetic and spatiotemporal data were processed using a custom-written MATLAB 2024a (Natick, MA) program. Force data were filtered using a low-pass fourth-order Butterworth filter with a cutoff frequency of 30 Hz. Heel-strike and toe-off events were identified using a 20 N threshold on the vertical ground reaction forces and these gait events determined each gait cycle. We visually inspected gait events to verify event detection and corrected if necessary. To evaluate propulsion and braking, we calculated peak anterior (AGRF) and posterior (PGRF, Appendix A) ground reaction forces, impulses and net anterior-posterior impulse (Appendix A). The speed of each belt was recorded for each trial and averaged over the duration of each trial. We calculated the position of participants’ COM to determine their position on the treadmill. To understand the spatiotemporal components of walking, we calculated stride length, stride time, step length and step time using force data (Appendix A). Kinetic measures were normalized to body weight and spatial measures were normalized to height. Each metric was averaged per limb for all gait cycles in each trial. To evaluate symmetry between limbs, asymmetry for any metric X was calculated as:

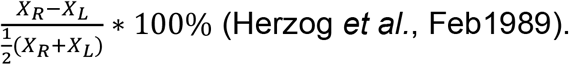

To explore the trends of the adaptive split-belt treadmill, we also ran mathematical simulations of speed using experimental data. A mathematical model of the sATM was applied to anterior ground reaction force (AGRF) data collected from the Normal Tied condition for a representative participant. We varied COM position relative to the treadmill center (0, 0.1, 0.2 m) with no changes in the AGRF data, as well as AGRF data (net impulse + 0.001, 0.005, 0.01 BW*s) with constant COM position relative to the treadmill center (at 0 m) to observe trends in the simulated velocity of the treadmill.

### 2.1.4 Statistical analysis

An a priori power analysis in RStudio 2024 (Posit, Boston, MA) for an equivalence test for propulsive impulse (bounds = 0.05, power = 0.8, std = 0.05) indicated 10 participants. Shapiro-Wilk tests were used to determine normality for all data.

To establish equivalence, we first looked at agreement between belts. We evaluated only the Split treadmill for walking speed as the belt speeds are always equal on the Tied treadmill. We evaluated the linear fit between the right and left belt. We then modeled the difference score in terms of condition and participant. Confidence intervals (CIs) for the difference score between trials were calculated using Tukey-HSD tests in JMP Pro (JMP^®^, Version *18*.*0*.*1* SAS Institute Inc., Cary, NC).

For all other measures, we looked at both Split and Tied treadmills. We examined the similarity between treadmills for all measures using repeated-measures two-way ANOVAs for each belt with factors of treadmill and condition. We also evaluated limb symmetry for all kinetic and spatiotemporal measures with repeated-measures two-way ANOVA with factors of treadmill and condition. CIs for the symmetry scores were calculated for each trial.

To determine differences between the belts for the Split Uneven condition, we ran paired Wilcoxon tests for all measures. All analyses where not specified were performed in Python (Vallat, 2018). Significance was set at 0.05 for all statistical analyses. Multiple comparisons were corrected using Bonferroni corrections.

## 3. Results

Participants’ self-selected walking (belt) speed was similar between Tied and Split treadmills (Table 1; Table 2; Fig 2A). The speeds of both belts showed linear agreement on the Split treadmill (adj. R^2^ = 0.409), where the difference in belt speeds was not significantly different than zero. Both belt speeds were significantly higher in the Hard (higher gamma) condition (Table 2). For the Split Uneven condition, belt speed did not show a significant difference between belts (p = 0.104).

**Table 1.** Kinetic metrics and COM position (Mean ± SD) for all walking trials.

| Treadmill | Tied |  | Split |  | Split Uneven |  |
| --- | --- | --- | --- | --- | --- | --- |
|  | Normal | Hard | Normal | Hard | Targeted Side | Non-targeted Side |
| **Belt Speed (m/s)** | R: 1.24±0.16<br>L: 1.24±0.16 | R: 1.3±0.16<br>L: 1.3±0.16 | R: 1.22±0.22<br>L: 1.26±0.18 | R: 1.28±0.22<br>L: 1.38±0.18 | 1.28±0.23 | 1.41±0.21 |
| **Peak AGRF (%BW)** | R: 20.91±3.81<br>L: 20.97±4.64 | R: 22.62±4.81<br>L: 22.67±5.45 | R: 19.61±5.33<br>L: 20.84±4.88 | R: 20.87±5.77<br>L: 22.67±5.38 | 21.65±5.77 | 22.05±5.17 |
| **AGRF Impulse (%BW\*s)** | R: 3.20±0.49<br>L: 3.13±0.51 | R: 3.33±0.62<br>L: 3.29±0.55 | R: 2.91±0.69<br>L: 3.07±0.58 | R: 2.95±0.69<br>L: 3.16±0.56 | 3.09±0.71 | 3.05±0.50 |
| **Center of Mass (m)** | R: 0.57±0.019<br>L: 0.57±0.019 | R: 0.79±0.009<br>L: 0.79±0.009 | R: 0.62±0.013<br>L: 0.62±0.013 | R: 0.65±0.046<br>L: 0.65±0.045 | 0.64±0.014 | 0.64±0.014 |

**Table 2.**
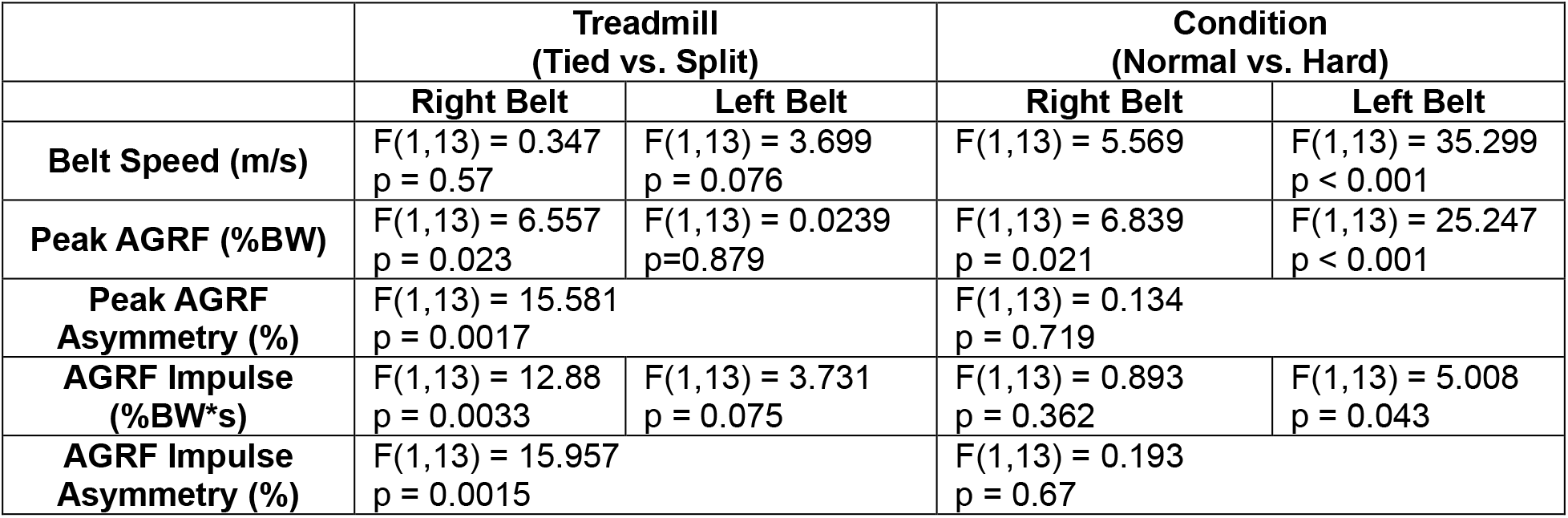
Statistical analysis results for Normal and Hard trials.

|  | Treadmill<br>(Tied vs. Split) |  | Condition<br>(Normal vs. Hard) |  |
| --- | --- | --- | --- | --- |
|  | Right Belt | Left Belt | Right Belt | Left Belt |
| **Belt Speed (m/s)** | F(1,13) = 0.347<br>p = 0.57 | F(1,13) = 3.699<br>p = 0.076 | F(1,13) = 5.569 | F(1,13) = 35.299<br>p < 0.001 |
| **Peak AGRF (%BW)** | F(1,13) = 6.557<br>p = 0.023 | F(1,13) = 0.0239<br>p = 0.879 | F(1,13) = 6.839<br>p = 0.021 | F(1,13) = 25.247<br>p < 0.001 |
| **Peak AGRF Asymmetry (%)** | F(1,13) = 15.581<br>p = 0.0017 |  | F(1,13) = 0.134<br>p = 0.719 |  |
| **AGRF Impulse (%BW\*s)** | F(1,13) = 12.88<br>p = 0.0033 | F(1,13) = 3.731<br>p = 0.075 | F(1,13) = 0.893<br>p = 0.362 | F(1,13) = 5.008<br>p = 0.043 |
| **AGRF Impulse Asymmetry (%)** | F(1,13) = 15.957<br>p = 0.0015 |  | F(1,13) = 0.193<br>p = 0.67 |  |

**Fig 2.**
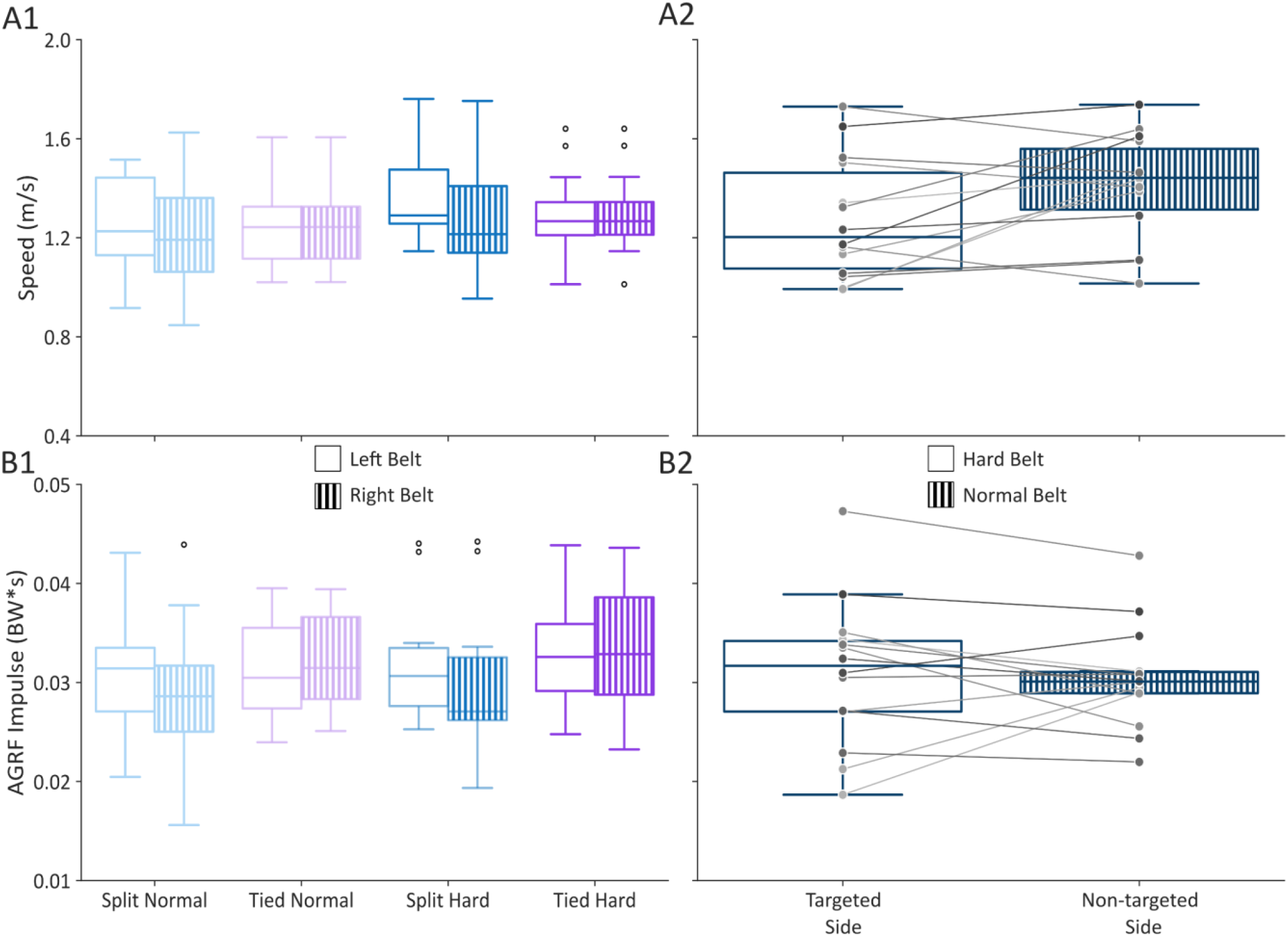
(A1) Walking Speed (B1) AGRF Impulse. Box plots represent data from all participants for the right and left belt for Normal and Hard conditions. (A2) Walking Speed (B2) AGRF Impulse. Box plots represent data from all participants for the belt with normal propulsion demand (Non-targeted) and increased propulsion demand (Targeted) for the Split Uneven condition.

Average AGRF impulse on the right belt was significantly higher on the Tied treadmill (Fig 2B; Table 2) but showed no significant effect of condition. For the left belt, AGRF impulse was higher in the Hard condition (higher gamma) but had no significant effect of treadmill (Table 2). Peak AGRF was significantly higher in the Hard condition for both belts and significantly higher in the Tied treadmill for the right belt but not the left belt (Table 2). AGRF impulse symmetry showed a statistically significant effect of treadmill but not condition (Table 2). Peak AGRF symmetry showed a significant effect of treadmill but not condition (Table 2). For the Split Uneven condition, AGRF impulse (p=0.583) and peak AGRF (p=0.855) did not show a significant difference between belts. In the few cases where the residual normality assumption for the ANOVAs was not met, the recovery action of outlier removal did not alter which effects were significant.

The COM position was equivalent between right and left strides (adj. R^2^ = 0.99) There was a significant interaction between treadmill and condition on COM position (F(1,13) = 250.184, p < 0.001). Participants’ COM was significantly more forward in the Hard condition for both Tied (Bonferroni-corrected p = 0.0027) and Split (Bonferroni-corrected p = 0.027).

Survey data were compared between all five conditions (Table 3). Participants reported that it was slightly easier to achieve and maintain speed and felt more comfortable and stable on the tied ATM. Participants also reported that it was easier to achieve and maintain speed and felt more comfortable and stable in the Normal condition than the Hard condition for both treadmills as well as the Split Uneven condition.

**Table 3.** Survey Data (Mean ± SD) for all walking trials.

| Treadmill | Tied |  | Split |  |  |
| --- | --- | --- | --- | --- | --- |
| Condition | Normal | Hard | Normal | Hard | Uneven |
| **Achieve Speed** | 6.5±0.65 | 4.21±1.48 | 5.21±1.31 | 4.71±1.49 | 4.43±1.65 |
| **Maintain Speed** | 6.43±0.51 | 4.86±1.56 | 4.71±1.38 | 4.86±1.17 | 4.21±1.37 |
| **Stability** | 6.64±0.5 | 5.36±1.39 | 5.07±1.49 | 5.14±1.29 | 4.57±1.79 |
| **Comfort** | 6.86±0.36 | 5.43±1.28 | 4.93±1.44 | 5.29±1.14 | 4.36±1.55 |

The simulated belt speed varied with variations in COM position as well as net anterior-posterior impulse (Fig 3). At the center of the treadmill (*COM*_*pos*_ = 0) with the original impulse data from a representative participant (experimental *v*_*PI*_), the simulated speed declined with the number of strides (Fig 3A). This indicates that the impulse was lower than required to maintain a speed of 1m/s at that position, and the participant was likely slightly closer to the front during the experiment, as moving forward on the treadmill would increase speed. Therefore, increasing the COM position relative to the treadmill center (increasing *COM*_*pos*_) corresponded to an increased simulated belt speed. Similarly, an increase in net impulse (increase in *v*_*PI*_) showed an increase in belt speed when COM position was held constant at the treadmill center (*COM*_*pos*_ = 0).

**Fig 3.**
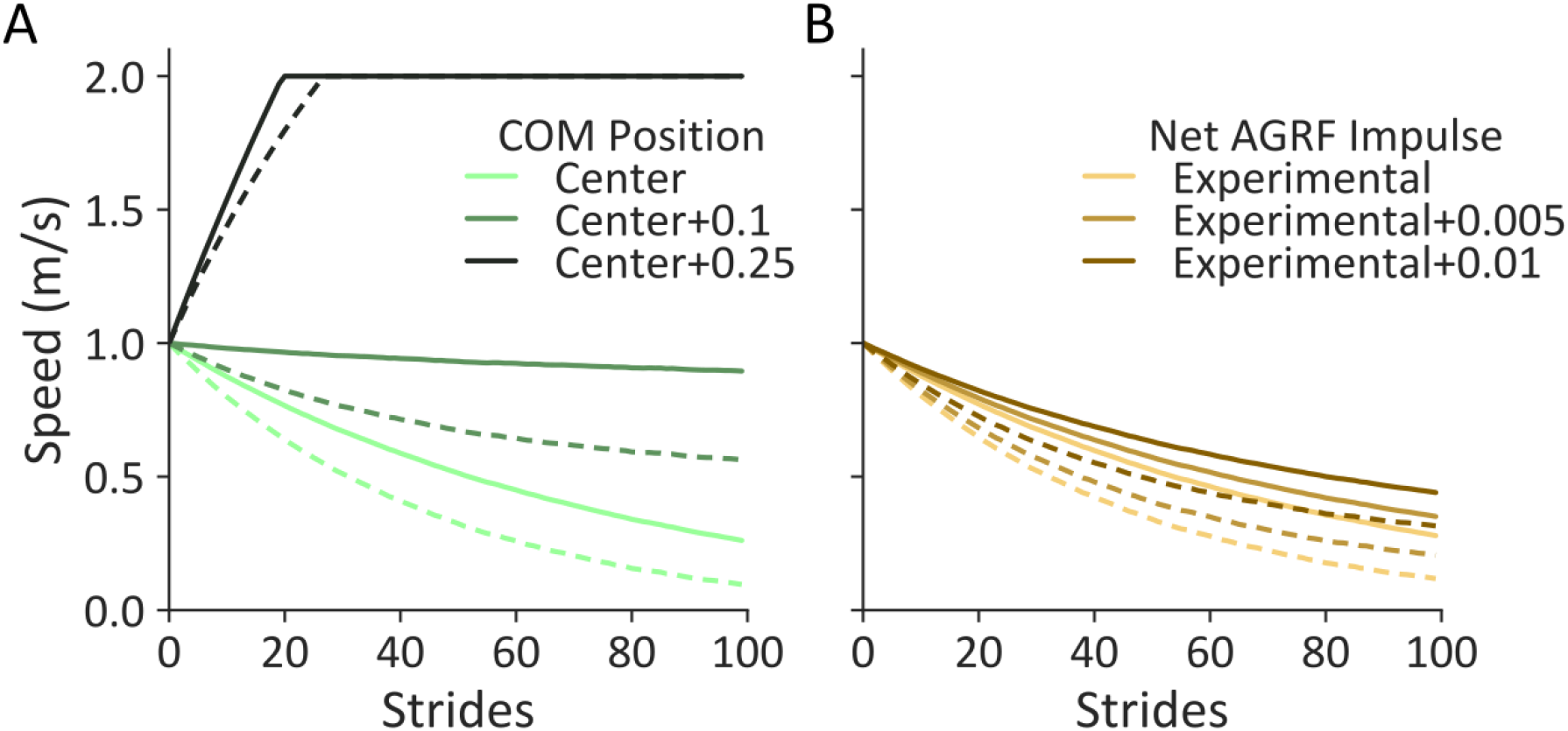
Simulation Results (A) Varying Position. Belt speed was simulated using the control equation for different center of mass (COM) positions relative to the center of the treadmill with the net AGRF impulse of a representative participant. (B) Varying Net Impulse. Belt speed was simulated using the control equation by varying the net AGRF impulse (by incrementing experimental data) of a representative participant. Solid lines depict the Split Normal condition, and dashed lines depict the Split Hard condition.

## 4. Discussion

We developed and validated a novel, adaptive split-belt treadmill. We hypothesized that propulsion and walking speed on the novel sATM would be equivalent to a previously developed tied adaptive treadmill, and further that the sATM could be preferentially weighted to encourage higher propulsion unilaterally while maintaining walking speed. Our findings partially support our hypothesis – participants maintained similar propulsion and walking speed between treadmills and belts. Interestingly, while most participants maintained similar walking speeds between belts when the sATM was preferentially unilaterally weighted, there was no significant difference between belts for propulsion on average. However, a few participants increased unilateral propulsion to maintain similar walking speeds, while others showed no change in propulsion with slight differences in walking speed. These results suggest that the response to unilateral weighting was user-specific, and the sATM could be tuned to individual responses.

We found that participants maintained their walking speed between Tied and Split treadmills. Post-hoc analysis showed that differences in speed between treadmills (Split - Tied) for Normal (Right: [−0.099,0.069], Left: [−0.078,0.125]) and Hard (Right: [−0.109,0.059], Left: [−0.021,0.183]) were within 95% CIs of differences in walking speed during continuous walking ([−0.127, 0.147]) obtained from prior literature (Paterson *et al*., 2008).

Participants also maintained similar speeds between belts on the Split treadmill. It is possible that participants did not perceive any differences between belts during the trials. 61% (17/28) of Split trials were in the healthy gait asymmetry range (*AGRF*_*asymm*_<=5%), suggesting that the sATM controller on average used a *v*_*PI*_ averaged over both limbs. Furthermore, previous work shows that healthy participants perceive belt speed differences during split-belt treadmill walking after a certain threshold of difference (95% CI: [0.025, 0.105]) is achieved (Hoogkamer *et al*., 2015). Post-hoc analysis of the perception threshold (absolute belt speed difference/sum of belt speeds) showed that the confidence interval of the perception threshold for all participants in both Split Normal ([0.033, 0.093]) and Split Hard (higher gamma, [0.035, 0.078]) conditions were within the aforementioned confidence intervals.

Participants had a higher self-selected speed in the Hard conditions (higher gamma), compared to the Normal conditions. While participants did not increase their AGRF impulse consistently between conditions, they were significantly closer to the front on the treadmill in the Hard condition for both treadmills. This suggests that participants modulated their position on the treadmill instead of AGRF impulse. Additionally, participants were instructed to walk at a comfortable speed, not their overground speed, and may not have sensed an “uncomfortable” change in walking speed between the Hard and Normal conditions.

Similar to walking speeds, propulsion (peak AGRF) was also consistent between treadmills and belts in the Normal and Hard conditions. While peak AGRF symmetry and AGRF impulse symmetry showed a significant effect of treadmill, symmetries for each condition were within healthy gait symmetry levels (±15.6, ±18.4 respectively) (Herzog *et al*., Feb1989). Therefore, we believe these bilateral differences are not clinically meaningful.

Participants increased their peak AGRF for the Hard conditions on both treadmills, suggesting that they were pushing off harder. These results indicate that by increasing the propulsion gain, we can encourage volitional bilateral changes in propulsion on the sATM. Peak AGRF is a good indicator of propulsion during walking in individuals post-stroke (Hsiao *et al*., 2016), therefore our results demonstrate a potential tool to encourage increased propulsion in stroke survivors.

Mathematically, we can demonstrate that increased propulsion is required for an increased gamma (increased propulsion demand), similar to previous Tied ATM work (Pariser *et al*., 2022). If we assume that users maintain similar positions on the treadmill at each stride, the change in speed between strides (*v*_*i*+1_ − *v*_*i*_) is driven primarily by the difference between the intermediate speed due to propulsion and the penalty factor based on the current belt speed 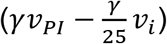. Therefore, under such conditions, to maintain the exact same speed on the treadmill regardless of gamma, the participant would need to maintain 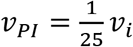. For human walking, the operational range of the treadmill lies within typical values of *v*_*PI*_ (net impulse) and belt speed from participants, which lead to 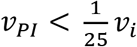; thus driving a slow decay of the belt speed, which is then regulated by the propulsion demand term (gamma). Therefore, for a higher gamma, the speed decay would be faster – requiring higher intermediate speed due to propulsion (*v*_*PI*_) to maintain speed. While the speed would increase faster if 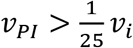 for a higher gamma, this condition lies outside the typical operational range, and participants do not maintain increased 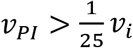 for more than a few gait cycles. Thus, this would only last until equilibrium is achieved, beyond which if the speed needs to be altered, the controller would once again fall under the previous conditions 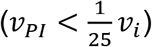 and require increased propulsion. Therefore, although increased propulsion might not be constantly applied throughout all strides in the Hard condition, participants would require consistently higher propulsion on average throughout the trial.

Despite different propulsion weights, the Split Uneven condition did not show significant differences in propulsion (peak AGRF and AGRF impulse) on the Targeted limb on average. However, on an individual level, a few participants increased their propulsion on the Targeted limb and achieved a smaller difference between belt speeds. Furthermore, net anterior-posterior impulse was higher on the Targeted limb and PGRF impulse were higher on the Non-targeted limb (higher braking) (Appendix A). Therefore, participants may have been able to maintain similar speeds on both belts by increasing their braking force on the Non-targeted limb and lowering the speed of that belt, while increasing the Targeted belt speed by increasing net impulse. Additionally, participants did not practice the Split Uneven condition and may have struggled to accommodate to the difference between belts. These results suggest that response to preferential weighting of unilateral propulsion is user-specific and may need to be tuned to individual responses to encourage a significant increase.

The simulation results demonstrate how the speed of each belt can be changed by modulating COM position or net anterior impulse. We have verified that the sATM controller algorithm behaves as anticipated by performing speed simulations using data collected from a participant. These results show how participants may have implemented different strategies to maintain comfortable walking speeds on the treadmill.

The survey results indicate that while participants perceived that achieving and maintaining their preferred speed was easier on the tied ATM as compared to the sATM for the Normal condition, it was equivalent between treadmills for the Hard conditions. Additionally, the Split Normal and Uneven conditions were similar to the Hard conditions in all survey responses. This suggests that while participants may have found it easier to use the tied ATM, they were able to control their walking speed on all sATM conditions with relative ease.

In the current work, users tended to modulate their position on the treadmill instead of propulsion to control the speed of the treadmill. Previous work has suggested that modifying the importance of position on the treadmill may result in changes in net impulse on the tied ATM (Downer *et al*., 2024). While drastically reducing the importance of position increased difficulty in achieving and maintaining walking speeds, it did not affect the comfort and stability of use of the tied ATM. Therefore, future work should explore modifying the COM term in the sATM controller to reduce reliance on the modulation of position on the treadmill while maintaining ease of use.

## 5. Conclusion

The present study demonstrates that walking speed can be modulated on an adaptive split-belt treadmill. In this work, we developed and validated an sATM that allows unilateral control strategies. Future work will explore different combinations of the propulsion gains on the sATM in the Split Uneven condition to determine gains that promote increased unilateral propulsion in healthy and post-stroke individuals.

## Supporting information

Data_WalkingSpeedModulatedOnSATM

SupplementalData_WalkingSpeedModulatedOnSATM

## 6. Acknowledgements

The authors were supported by the National Institute of Health (NIH #HD111071) and an Institutional Development Award (IDeA) from the National Institute of General Medical Sciences of the National Institute of Health under grant number U54-GM104941 (PI: Hicks)

## 8. Preprint

A preprint of this manuscript is available at https://doi.org/10.1101/2025.06.03.657157.

## 9. Data Statement

The data will be made available upon request.

## APPENDIX A

We analyzed secondary kinetic and spatiotemporal measures to provide further insight into participants’ walking on both treadmills for all conditions (Table A1).

Average PGRF impulse showed a significant effect of treadmill (Right: F(1,13), p = 0.0012; Left: F(1,13), p = 0.0093) and condition (Right: F(1,13), p = 0.020; Left: F(1,13), p = 0.0031) for both belts. PGRF impulse symmetry showed no significant effect of treadmill or condition. Peak PGRF had a significant effect of condition for both belts (F(1,13), p < 0.001) and treadmill for the right belt (F(1,13), p = 0.043). There was no significant effect of treadmill or condition on peak PGRF symmetry. Net anterior-posterior impulse on the left belt showed a significant effect of treadmill (F(1,13), p = 0.028) and condition (F(1,13), p = 0.027), while the right belt had no significant differences. Net anterior-posterior impulse symmetry showed no significant differences between treadmill or condition.

AGRF time showed a significant effect of treadmill (F(1,13), p = 0.033) and condition (F(1,13), p = 0.042) on the left belt. PGRF time showed a significant effect of condition on both belts (Right: F(1,13), p = 0.0031; Left: F(1,13), p < 0.001) and treadmill on the left belt (F(1,13), p = 0.027) but not the right. There was no significant effect of treadmill or condition on AGRF time symmetry and PGRF time symmetry. The peak AGRF time was significantly different between conditions for both belts (Right: F(1,13), p = 0.0096; Left: F(1,13), p < 0.001) and between treadmills for the left belt (F(1,13), p =0.013) but not the right. There was no significant effect of treadmill or condition on the peak AGRF time symmetry. The peak braking time was significantly different between conditions for both belts (Right: F(1,13), p = 0.038; Left: F(1,13), p = 0.0041) and between treadmills for the left belt (F(1,13), p = 0.011) but not right. There was a significant effect of treadmill (F(1,13), p = 0.0038) but not condition on the symmetry of the time of peak braking.

Stride length showed a significant effect of condition on the left belt (F(1,13), p = 0.002). Stride length symmetry showed no significant differences. Step length had a significant effect of condition on the right belt (F(1,13), p < 0.001) and effect of treadmill on the left belt (F(1,13), p = 0.043). There was a significant effect of treadmill on step length symmetry (F(1,13), p = 0.019). Stride time showed a significant effect of condition on both belts (Right: F(1,13), p = 0.0054; Left: F(1,13), p = 0.0049). Stride time symmetry showed no significant differences. Step time was significantly different between conditions for both belts (Right: F(1,13), p = 0.0038; Left: F(1,13), p = 0.029) and between treadmills for the right belt (F(1,13), p = 0.014). Step time symmetry showed no significant effect of treadmill or condition.

For Split Uneven, there was no significant difference between belts for PGRF time, peak PGRF time, stride length, stride time, step length, or step time. PGRF impulse (F(1,13), p = 0.024) and peak PGRF (F(1,13), p = 0.049) was greater for the Non-targeted limb. Net anterior-posterior impulse (F(1,13), p = 0.029), AGRF time (F(1,13), p = 0.0012), and peak AGRF time (F(1,13), p = 0.025) were greater for the Targeted limb.

**Table A1.** Secondary kinetic and spatiotemporal metrics (Mean ± SD).

| Treadmill | Tied |  | Split |  | Split Uneven |  |
| --- | --- | --- | --- | --- | --- | --- |
|  | Normal | Hard | Normal | Hard | Non-Targeted Side | Targeted Side |
| **Peak PGRF (%BW)** | R: 19.15±4.95<br>L: 19.62±4.57 | R: 21.63±5.74<br>L: 22.34±5.39 | R: 18.19±5.33<br>L: 18.93±4.74 | R: 19.86±5.42<br>L: 21.44±5.12 | 22.21±6.21 | 20.75±5.81 |
| **PGRF Impulse (%BW\*s)** | R: 3.29±0.53<br>L: 3.37±0.50 | R: 3.57±0.57<br>L: 3.66±0.61 | R: 3.08±0.65<br>L: 3.19±0.55 | R: 3.13±0.54<br>L: 3.38±0.47 | 3.48±0.63 | 3.22±0.56 |
| **Net Impulse (%BW)** | R: -0.09±0.25<br>L: -0.24±0.25 | R: -0.24±0.27<br>L: -0.36±0.24 | R: -0.18±0.21<br>L: -0.11±0.24 | R: -0.18±0.44<br>L: -0.22±0.27 | -0.43±0.27 | -0.12±0.22 |
| **AGRF Time (s)** | R: 0.32±0.026<br>L: 0.32±0.030 | R: 0.32±0.030<br>L: 0.31±0.027 | R: 0.31±0.031<br>L: 0.31±0.019 | R: 0.31±0.022<br>L: 0.30±0.020 | 0.29±0.020 | 0.31±0.026 |
| **PGRF Time (s)** | R: 0.39±0.037<br>L: 0.39±0.031 | R: 0.37±0.037<br>L: 0.38±0.030 | R: 0.38±0.041<br>L: 0.38±0.032 | R: 0.37±0.043<br>L: 0.36±0.034 | 0.36±0.038 | 0.37±0.036 |
| **Peak AGRF Time (s)** | R: 0.61±0.05<br>L: 0.62±0.05 | R: 0.59±0.04<br>L: 0.60±0.04 | R: 0.60±0.05<br>L: 0.59±0.04 | R: 0.58±0.05<br>L: 0.57±0.05 | 0.56±0.048 | 0.59±0.51 |
| **Peak PGRF Time (s)** | R: 0.13±0.02<br>L: 0.13±0.03 | R: 0.13±0.02<br>L: 0.12±0.02 | R: 0.13±0.02<br>L: 0.12±0.02 | R: 0.12±0.02<br>L: 0.11±0.02 | 0.11±0.025 | 0.12±0.024 |
| **Stride Length** | R: 0.48±0.061<br>L: 0.48±0.061 | R: 0.49±0.058<br>L: 0.49±0.057 | R: 0.46±0.079<br>L: 0.47±0.067 | R: 0.46±0.068<br>L: 0.50±0.069 | 0.87±0.11 | 0.79±0.13 |
| **Step Length** | R: 0.24±0.031<br>L: 0.24±0.030 | R: 0.24±0.029<br>L: 0.25±0.029 | R: 0.23±0.032<br>L: 0.23±0.037 | R: 0.25±0.033<br>L: 0.24±0.032 | 0.40±0.058 | 0.42±0.051 |
| **Stride Time (s)** | R: 1.12±0.07<br>L: 1.12±0.07 | R: 1.09±0.06<br>L: 1.09±0.06 | R: 1.09±0.07<br>L: 1.09±0.07 | R: 1.06±0.06<br>L: 1.06±0.06 | 1.06±0.063 | 1.06±0.062 |
| **Step Time (s)** | R: 0.56±0.03<br>L: 0.56±0.04 | R: 0.54±0.03<br>L: 0.55±0.03 | R: 0.54±0.03<br>L: 0.55±0.04 | R: 0.52±0.03<br>L: 0.54±0.04 | 0.54±0.036 | 0.52±0.038 |

## References

Awad, L.N. et al. (2020) “These legs were made for propulsion: advancing the diagnosis and treatment of post-stroke propulsion deficits,” Journal of NeuroEngineering and Rehabilitation, 17(1), p. 139. Available at: 10.1186/s12984-020-00747-6.

Dickstein, R. (2008) “Rehabilitation of Gait Speed After Stroke: A Critical Review of Intervention Approaches,” Neurorehabilitation and Neural Repair, 22(6), pp. 649–660. Available at: 10.1177/1545968308315997.

Donlin, M.C. et al. (2022) “Adaptive treadmill walking encourages persistent propulsion,” Gait & Posture, 93, pp. 246–251. Available at: 10.1016/j.gaitpost.2022.02.017.

Downer, K.E. et al. (2024) “How Important is Position in Adaptive Treadmill Control?,” Journal of Biomechanical Engineering, 146(1), p. 011006. Available at: 10.1115/1.4063823.

Duncan, P.W. et al. (2011) “Body-Weight–Supported Treadmill Rehabilitation after Stroke,” New England Journal of Medicine, 364(21), pp. 2026–2036. Available at: 10.1056/NEJMoa1010790.

Fritz, S. and Lusardi, M. (2009) “White Paper: ‘Walking Speed: the Sixth Vital Sign’:,” Journal of Geriatric Physical Therapy, 32(2), pp. 2–5. Available at: 10.1519/00139143-200932020-00002.

Gelaw, A.Y. et al. (2019) “Effectiveness of treadmill assisted gait training in stroke survivors: A systematic review and meta-analysis,” Global Epidemiology, 1, p. 100012. Available at: 10.1016/j.gloepi.2019.100012.

Herzog, W. et al. (Feb 1989) “Asymmetries in ground reaction force patterns in normal human gait,” Medicine and Science in Sports and Exercise, 21(1), pp. 110–114. Available at: 10.1249/00005768-198902000-00020.

Hoogkamer, W. et al. (2015) “Gait asymmetry during early split-belt walking is related to perception of belt speed difference,” Journal of Neurophysiology, 114(3), pp. 1705–1712. Available at: 10.1152/jn.00937.2014.

Hsiao, H. et al. (2015) “The relative contribution of ankle moment and trailing limb angle to propulsive force during gait,” Human Movement Science, 39, pp. 212–221. Available at: 10.1016/j.humov.2014.11.008.

Hsiao, H. et al. (2016) “Evaluation of measurements of propulsion used to reflect changes in walking speed in individuals poststroke,” Journal of Biomechanics, 49(16), pp. 4107–4112. Available at: 10.1016/j.jbiomech.2016.10.003.

Kempski, K.M. et al. (2019) “Dynamic structure of variability in joint angles and center of mass position during user-driven treadmill walking,” Gait & Posture, 71, pp. 241–244. Available at: 10.1016/j.gaitpost.2019.04.031.

Lauzière, S. et al. (2014) “Understanding Spatial and Temporal Gait Asymmetries in Individuals Post Stroke,” International Journal of Physical Medicine & Rehabilitation, 02(03). Available at: 10.4172/2329-9096.1000201.

Pariser, K.M. et al. (2022) “Adaptive treadmill control can be manipulated to increase propulsive impulse while maintaining walking speed,” Journal of Biomechanics, 133, p. 110971. Available at: 10.1016/j.jbiomech.2022.110971.

Paterson, K.L. et al. (2008) “The Reliability of Spatiotemporal Gait Data for Young and Older Women During Continuous Overground Walking,” Archives of Physical Medicine and Rehabilitation, 89(12), pp. 2360–2365. Available at: 10.1016/j.apmr.2008.06.018.

Patterson, S.L. et al. (2007) “Determinants of Walking Function After Stroke: Differences by Deficit Severity,” Archives of Physical Medicine and Rehabilitation, 88(1), pp. 115–119. Available at: 10.1016/j.apmr.2006.10.025.

Ray, N.T., Knarr, B.A. and Higginson, J.S. (2018) “Walking speed changes in response to novel user-driven treadmill control,” Journal of Biomechanics, 78, pp. 143–149. Available at: 10.1016/j.jbiomech.2018.07.035.

Ray, N.T., Reisman, D.S. and Higginson, J.S. (2020) “Walking speed changes in response to user-driven treadmill control after stroke,” Journal of Biomechanics, 101, p. 109643. Available at: 10.1016/j.jbiomech.2020.109643.

Reisman, D.S. et al. (2007) “Locomotor adaptation on a split-belt treadmill can improve walking symmetry post-stroke,” Brain, 130(7), pp. 1861–1872. Available at: 10.1093/brain/awm035.

Reisman, D.S. et al. (2009) “Split-Belt Treadmill Adaptation Transfers to Overground Walking in Persons Poststroke,” Neurorehabilitation and Neural Repair, 23(7), pp. 735–744. Available at: 10.1177/1545968309332880.

Reisman, D.S. et al. (2013) “Repeated Split-Belt Treadmill Training Improves Poststroke Step Length Asymmetry,” Neurorehabilitation and Neural Repair, 27(5), pp. 460–468. Available at: 10.1177/1545968312474118.

Sloot, L.H., Van Der Krogt, M.M. and Harlaar, J. (2014) “Self-paced versus fixed speed treadmill walking,” Gait & Posture, 39(1), pp. 478–484. Available at: 10.1016/j.gaitpost.2013.08.022.

Stergiou, N. and Decker, L.M. (2011) “Human movement variability, nonlinear dynamics, and pathology: Is there a connection?,” Human Movement Science, 30(5), pp. 869–888. Available at: 10.1016/j.humov.2011.06.002.

Vallat, R. (2018) “Pingouin: statistics in Python,” Journal of Open Source Software, 3(31), p. 1026. Available at: 10.21105/joss.01026.

